# Gene regulatory network (GRN) embedded agents connect cellular decision making to human pluripotent stem cell derived germ layer-like pattern formation

**DOI:** 10.1101/2020.10.06.327650

**Authors:** Himanshu Kaul, Nicolas Werschler, Mukul Tewary, Andrew Hagner, Joel Ostblom, Daniel Aguilar-Hidalgo, Peter W. Zandstra

## Abstract

The emergence of germ layers in embryos during gastrulation is a key developmental milestone. How morphogenetic signals engage the regulatory networks responsible for early embryonic tissue patterning is incompletely understood. To understand this, we developed a gene regulatory network (GRN) model of human pluripotent stem cell (hPSC) lineage commitment and embedded it into ‘cellular’ agents that respond to a dynamic signalling microenvironment. We found that cellular pattern order, composition, and dynamics were predictably manipulable based on the GRN wiring. We showed that feedback between OCT4, and BMP and WNT pathways created a dynamic OCT4 front that mediates the spatiotemporal evolution of developmental patterns. Translocation of this radial front can be predictively disrupted in vitro to control germ-layer pattern composition. This work links the emergence of multicellular patterns to regulatory network activity in individual hPSCs. We anticipate our approach will help to understand how GRN structure regulates organogenesis in different contexts.

## INTRODUCTION

Gastrulation is the phenomenon that allocates the seemingly equivalent epiblast cells to the three germ layers: ectoderm, mesoderm, and endoderm. These germ layers contribute to the formation of all tissues and organs. The mechanisms underlying germ layer emergence in humans remain elusive due to technical and ethical challenges. Seeding human pluripotent stem cell (hPSC) colonies on extracellular matrix micropatterns of defined shape and size provides a model to study elements of this phenomenon (Warmflash *et al*., 2014; Tewary *et al*., 2017). Upon stimulation by Bone Morphogenetic Protein-4 (BMP4), micropatterned hPSC colonies selforganise with SOX2+ (ectoderm-associated), Brachyury+ (mesoderm-associated), SOX17+ (endoderm-associated), and CDX2+ (extraembryonic-associated) cells that are radially segregated. This pattern has been explained using mathematical reaction-diffusion (RD) models based on BMP4 activation and autoregulation (Etoc *et al*., 2016; Tewary *et al*., 2017). The patterning signaling cascade has been extended to include a BMP → Wingless Integrated (WNT) → NODAL pathway hierarchy in hPSCs (Martyn *et al*., 2018). Evidence exists that this cascade evolves dynamically in space and time yielding neighbouring domains in a sequential manner (Chhabra *et al*., 2019; Tewary *et al*., 2019).

Despite advances in our understanding of morphogen signaling dynamics, how these exogenous signals engage the PSC fate decision-making regulatory networks during germ-layer fate commitment remains incompletely understood. Conventionally, this decision-making has been studied as an interplay between single cell heterogeneity and gene regulation. In this view differentiation cues amplify the cells’ initial lineage bias until their expression profile changes to reflect a new, stable cell type or expression state (Semrau and van Oudenaarden, 2015). Cell type, thus, is an emergent property of the underlying gene regulatory network (GRN) (Semrau and van Oudenaarden, 2015).

Pluripotency-maintaining GRNs in “individual” mouse and human PSCs have been extensively studied by us and others (Yachie-Kinoshita *et al*., 2018; Dunn *et al*., 2014; Dunn *et al*., 2019; Lin *et al*., 2018; Xu *et al*., 2014; Okawa and del Sol, 2015; Ng and Surani, 2011; Kalmar *et al*., 2009). Briefly, core pluripotency factors OCT4, NANOG, and SOX2 are closely linked via autoregulatory interactions (Boyer *et al*., 2005; Yan *et al*., 2013). Systematic knockout studies have revealed that while OCT4 & NANOG are co-regulated, SOX2 can be manipulated independently (Wang *et al*., 2012). Additionally, OCT4 expression is supported by β-catenin, the transcriptional effector of WNT3, as discovered via GSK3 antagonism (Sato *et al*., 2004; Sokol, 2011), likely dependent on WNT signaling duration and dosage (Davidson *et al*., 2012). Further, alternative splicing that enhances the DNA binding affinity of the forkhead family transcription factor FOXP1 has been shown to stimulate OCT4 and NANOG gene expression (Gabut *et al*., 2011). The pluripotency factors also mediate lineage commitment associated transcription factors (TFs) (Wang *et al*., 2012). Key here is the role of OCT4 expression in mediating differentiation to mesendoderm (when “high”) and extraembryonic fates (when “low”) upon exposure to BMP4. β-catenin and OCT4 interplay further drives primitive streak induction (Funa *et al*., 2015; Blauwkamp *et al*., 2012). Subsequently, Brachyury (TBXT) interacts with SMAD1 and SMAD2/3 signalling to regulate mesoderm and endoderm expression, respectively, by targeting a wide array of TFs, including CDX2, WNT3, EOMES, FOXA1 (Faial *et al*., 2015). Although this overview provides an abridged version of PSC regulatory control, both in terms of supporting components (Semrau and van Oudenaarden, 2015) and additional layers of control, it nevertheless serves as a sufficiently complex and predictive foundation for our tissue-patterning investigations.

Using abstracted GRN as a basis, cell fate responses to external ligands can be approximated and predicted by a relatively simple process of molecular computation using Boolean logic (Dunn *et al*., 2014; Dunn *et al*., 2019; Yachie-Kinoshita *et al*., 2018). For example, Dunn *et al*. (2014) used a data constrained computational approach to derive a set of functionally validated network components and interaction combinations to explain mouse PSC behavior. Yachie-Kinoshita *et al*. (2018) extended this strategy to early PSC differentiation in response to input signals. Such efforts have not yet been undertaken to explain germ layer emergence in human PSCs and, additionally, are limited to simulating the behavior of the individual cell in the absence of cellcell communication networks.

Herein, we bridge this gap and explore how contextualising the GRN topology with spatially evolving morphogens influence multicellular patterning. We used agent-based modelling (ABM) to investigate the multiscale interactions that govern self-organisation in hPSC colonies using methodological advantages that are described in-depth elsewhere (Kaul and Ventikos, 2015b). Briefly, multiple single cell PSC GRNs were discretised in space using agents (virtual cells) linked to morphogen gradients based on reaction-diffusion (virtual environment). Using this so-called GARMEN (<u>G</u>RN-<u>A</u>gent-<u>R</u>D <u>M</u>odelling <u>E</u>nvironment) platform, we show that spatial patterning is driven by a mutually reciprocal dynamic interplay between the GRN and local signal gradients. We show how single cell GRN wiring influences tissue-like pattern formation. We further demonstrate that WNT and BMP pathways guide the spatial position of OCT4 expressing cells in time, resulting in the emergence of a dynamic inward-translocating OCT4 front. Overall, our work links a minimal single hPSC GRN to higher order tissue-like patterns in a predictive manner, providing mechanistic insights into early fate control.

## RESULTS

### A minimal GRN captures exit from pluripotency to germ-layer fates in hPSCs

To identify the mechanisms that mediate germ-layer patterning in hPSCs, we first conducted an analysis of literature to identify genes, TFs, and signalling pathways implicated in pluripotency and germ layer emergence. As we intended to create a minimal GRN that captured both the maintenance of pluripotency and exit to germ-layer states, we focused on a narrow set of genes, signals, and interactions for model development. For example, we set-aside NANOG due to its lineage restricted role in maintaining pluripotency (Wang *et al*., 2012) and NODAL signalling given that BMP & WNT pathways have been shown to successfully trigger exit from pluripotency to germ-layer fates (Tewary *et al*., 2017; Chhabra *et al*., 2019; Martyn *et al*., 2018). Similarly, other germ-layer fate associated TFs, such as EOMES, FOXA1, OTX2, were also not directly considered at this time. Achieving this level of parsimony helped to keep the GRN (and the eventual multiscale model) tractable. The key components and interactions we relied on to compose the GRN are summarised in **Table S1** and directed graph of intracellular decisionmaking generated from these components in hPSCs is depicted in **Figure 1A**. This directed graph included pluripotency-associated genes OCT4 and SOX2, primitive streak-associated factors TBXT and SOX17, and extraembryonic lineage specifier CDX2. We also included key signalling pathways by linking gene states to signalling activities. Specifically, we worked with BMP4 and WNT3 activity, also including their inhibitors NOGGIN and DKK, respectively, in the model. To complete the loop (i.e. Signal→GRC→Signal), we built regulatory edges linking genes to signalling pathways (e.g. OCT4→WNT3). To transform this directed graph into a dynamical model, we adopted a multilevel logic formalism, to abstract GRN node states as a ternary variable (OFF, LOW, HIGH), an approach similar to Collombet *et al*. (2017). The regulatory relationships and logic are listed in **Table 1**. Governing interactions between the nodes of this multilevel model were manually interpreted following the review of the aforementioned literature and the interactions proposed therein (see **Methods** and **Table S1**).

**Figure 1.**
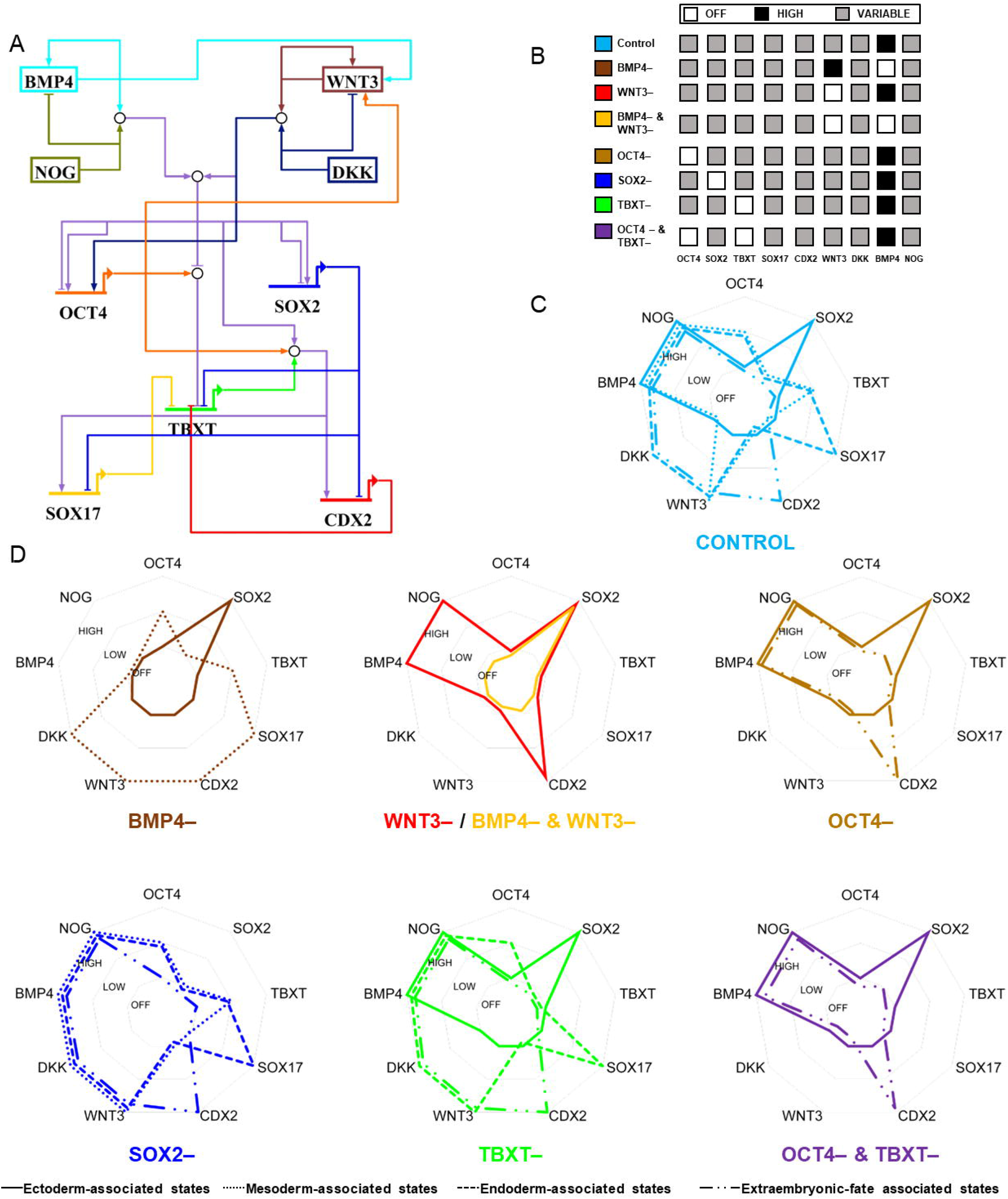
GRN capturing exit from pluripotency to germ-layer fates. A. The cell decision-making GRN. The hollow circles with converging and diverging arrows act as logic gates for this model where the relation between the input and output is based on AND, OR, or NOT logic. B. Inputs for the various conditions (control and perturbations) simulated by the GRN shown in A. The white square indicates the node will be continuously set to OFF, black square indicates the node will be continuously set to HIGH, and the grey square indicates that the node can assume any state between OFF, LOW, HIGH. C. D. Radar plots showing states of the GRN nodes observed in attractors (stable states and terminal complex SCCs) when conditions shown in B are simulated asynchronously. The plot represents the node states for various germ layer-associated states, as listed in Supplementary Table S4 (ectoderm-associated state in *solid line*; mesoderm-associated state in *round dots*; endoderm-associated state in *square dots*; and extraembryonic-associate state in *long dash dot dot*. Refer to Supplementary Data S1-2 for details regarding the nature of attractors yielded by the various conditions.

**Table 1.**
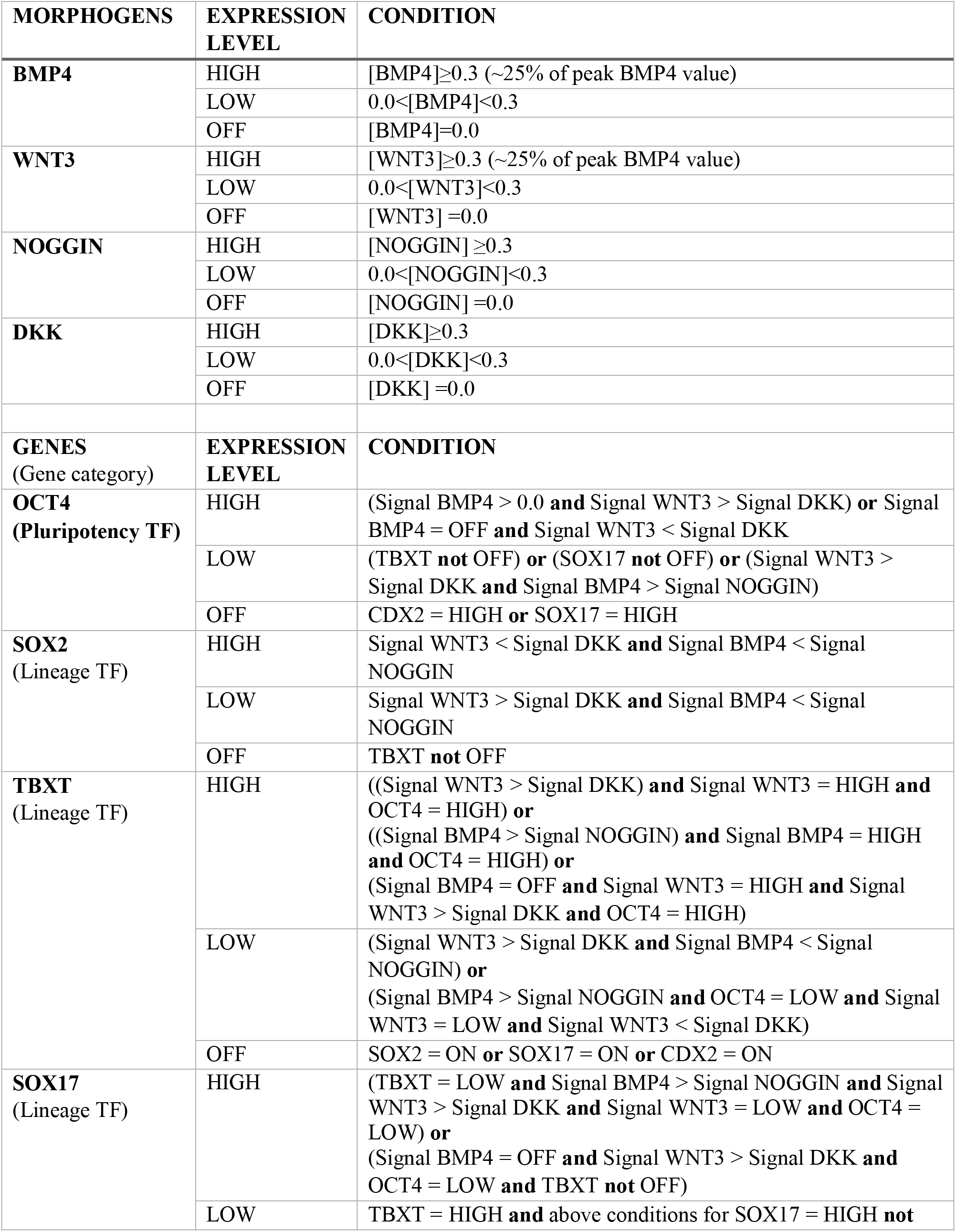
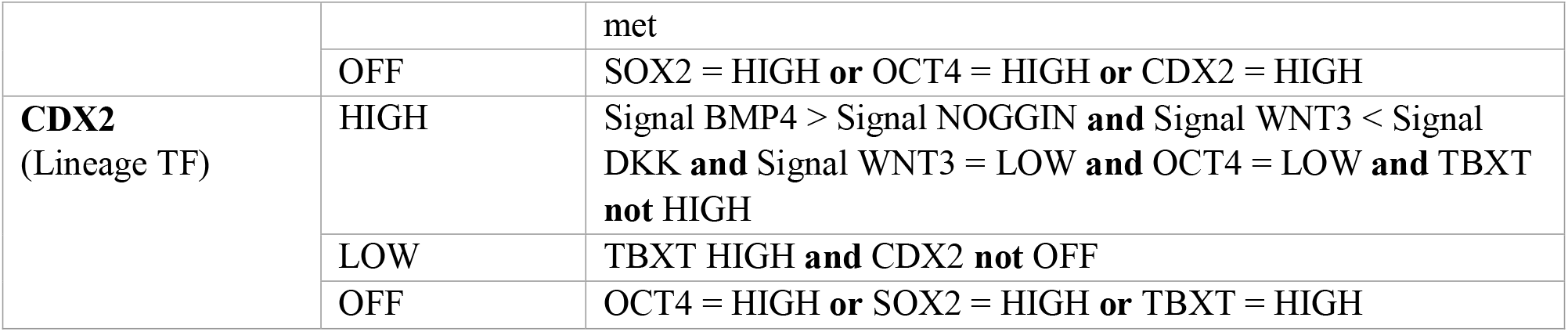
Gene regulatory network rules regulating differentiation related decision-making

We next assessed the ability of this GRN to approximate germ-layer fate commitment. We achieved this by simulating the model in the *Gene Interaction Network Simulation* (GINsim) platform (Naldi *et al*., 2009) and assessing whether 1) the resulting attractor states corresponded to germ-layer fates, i.e. we looked for both germ-layer associated unique steady states and terminal strongly connected components (SCCs), the latter defined as SCCs that do not reach any other components but themselves (Bérenguier *et al*., 2013), and 2) there exist unique paths from the initial pluripotent state to germ-layer states. Comparisons were made against existing literature, as tabulated in **Table S2**.

Following our experimental system (Tewary *et al*., 2017), we simulated the GRN responses after BMP4 addition (**Figure 1B**) and ran the model asynchronously (Yachie-Kinoshita *et al*., 2018). As initial conditions, OCT4 and SOX2 were turned HIGH, to mimic cellular pluripotency, BMP4 was continuously set to HIGH to match experimental conditions, whereas the levels of other nodes were randomly set (**Table S3**). This simulation yielded unique paths from the pluripotent to ectoderm−, mesoderm−, endoderm−, and extraembryonic-associated states. It also yielded a terminal complex SCC that consisted of 648 states (**Data S1**). This terminal complex SCC captured ectoderm−, mesoderm−, endoderm−, and extraembryonic-associated states (**Figure 1C and Data S2**). Together, these outputs suggested that if continually exposed to BMP4, hPSCs can access a number of germ-layer states.

We next simulated the response of this GRN (**Figure 1A**) to select perturbations (detailed in **Figure 1B** and initial conditions shown in **Table S3**). Perturbing the network entailed setting the activity level of the relevant components continuously to OFF (represented by ‘–’). Setting BMP4 to OFF and simulating the model by activating WNT3 (and keeping it continuously HIGH) resulted in two stable state attractors: one representing the ectoderm-associated state and the other capturing a mesendoderm-associated state (**Figure 1D and Data S1, S2**). This simulation also yielded unique paths from pluripotency to all germ layer-associated states. This suggested that the WNT pathway was sufficient to trigger germ-layer markers in hPSC colonies, as has been documented elsewhere (Martyn *et al*., 2018). Setting WNT3 activity levels to OFF (and triggering differentiation by setting BMP4 continuously to HIGH), on the other hand, led to a unique stable state with only SOX2 and CDX2 set to HIGH (besides WNT3 and DKK) (**Figure 1D**), as has been observed previously (Martyn *et al*., 2018). Setting activity levels of both BMP4 and WNT3 to OFF continuously led to a unique ectodermal steady state (**Figure 1D**). This condition reflected a ‘default’ state to which the system will return to in the absence of exogenous cues, consistent with experimental evidence that lack of external signals push PSCs to ectodermal fates (Blauwkamp *et al*., 2012; Fathi, Eisa-Beygi and Baharvand, 2017; Smukler *et al*., 2006).

Setting OCT4 levels continuously to OFF, on the other hand, resulted in loss of the mesendoderm state in the terminal complex SCC (**Figure 1D**), with unique paths only to ectoderm-associated and extraembryonic-associated states, consistent with an earlier investigation (Wang *et al*., 2012). On the contrary, setting SOX2 continuously OFF led to the loss of ectoderm-associated states and unique paths to mesoderm−, endoderm−, and extraembryonic-associated states, in agreement with published literature (Wang *et al*., 2012). Turning TBXT continuously OFF led to the loss of mesoderm states, but yielded ectoderm−, endoderm−, and extraembryonic-associated states, consistent with earlier findings (Faial *et al*., 2015). Finally, continuously turning OCT4 and TBXT OFF led to a terminal complex SCC with no mesoderm- and endoderm-associated fates, as has been reported previously (Faial *et al*., 2015; Wang *et al*., 2012).

To summarise, these results support the use our minimal GRN as a test system to provisionally capture germ layer-associated states from a pluripotent initial state given different input signaling conditions.

### Embedding GRNs within morphogen-sensing agents helps capture spatial and temporal emergence of germ layers

We next investigated whether this GRN could capture the spatial and temporal emergence of the germ-layer patterns in BMP-treated populations of hPSCs. We approached this by building a framework that allowed us to discretise gene networks explicitly in space and study interactions between these networks. Patterning has been previously simulated based on direct linkage between genes and signalling pathways via activator-inhibitor Turing-like mechanisms (Raspopovic *et al*., 2014; Cotterell and Sharpe, 2010; Cotterell, Robert-Moreno and Sharpe, 2015). We developed our framework instead by embedding our GRN in cellular “agents” (**Figure S1A**). Agents are computational systems that are situated within an environment and capable of engaging in flexible, autonomous action within that environment to meet their design objectives (Kaul and Ventikos, 2015b; Wooldridge, 1997; Jennings, 2000). They are a representational formalism for cells (Kaul and Ventikos, 2015b) and have been utilised as virtual cells in non-clinical and clinical contexts (Kaul, Cui and Ventikos, 2013; Kaul *et al*., 2015; Abbott, Forrest and Pienta, 2006; Segovia-Juarez, Ganguli and Kirschner, 2004; Casal *et al*., 2005; Bailey, Thorne and Peirce, 2007; Martin *et al*., 2017; Lee *et al*., 2019; Pothen, Poynter and Bates, 2015; Saunders *et al*., 2017; Saunders *et al*., 2019; Chachi *et al*., 2019; An, 2004; Adra *et al*., 2010; Coakley, Smallwood and Holcombe, 2006; Sun *et al*., 2009; Tao *et al*., 2007; Li *et al*., 2013).

We first created an agent-based model (ABM) integrating modules for the virtual cells to account for physical interactions, proliferation, morphogen release and sensing, and differentiation (based on the GRN). These virtual cells were assigned the following key characteristics: phenotype (combination of final GRN states, as outlined in **Table S4**), spatial coordinates, cell cycle state, and ability to detect biochemical gradients (**Figures S1B and S1C**). To this cell model we added our GRN as another module. The GRN regulated the signalling activity and differentiation state of the virtual cells (**Figure S1D**). This model allowed us to capture hPSC fate evolution based on the GRN-driven activity of individual cells.

To connect GRN activity to exogenous cues, we enabled the virtual cells to detect morphogens (**Figure S1E**) in their local microenvironment. This was achieved by coupling the *virtual cell with the GRN* paradigm to a four-component reaction-diffusion model that simulated the BMP and WNT pathways (refer to **Methods, Figure S1F**). This four-component model is referred to as the *RD module* of the platform henceforth. The RD→GRN→RD loop was closed by tying the activation of the RD module to the GRN via agents that had not transitioned to mesendoderm-associated states. Refer to *Methods* for more details, especially the coupling of various paradigms. The architecture of this GRN Agent-based Reaction-Diffusion Modelling Environment (GARMEN) is represented in **Figure 2A** and its implementation shown in **Figure S1G** and **BOX 1**. The in silico geometry where the computations were conducted and its discretisation is shown in **Figures S2A-S2C**.

**Figure 2.**
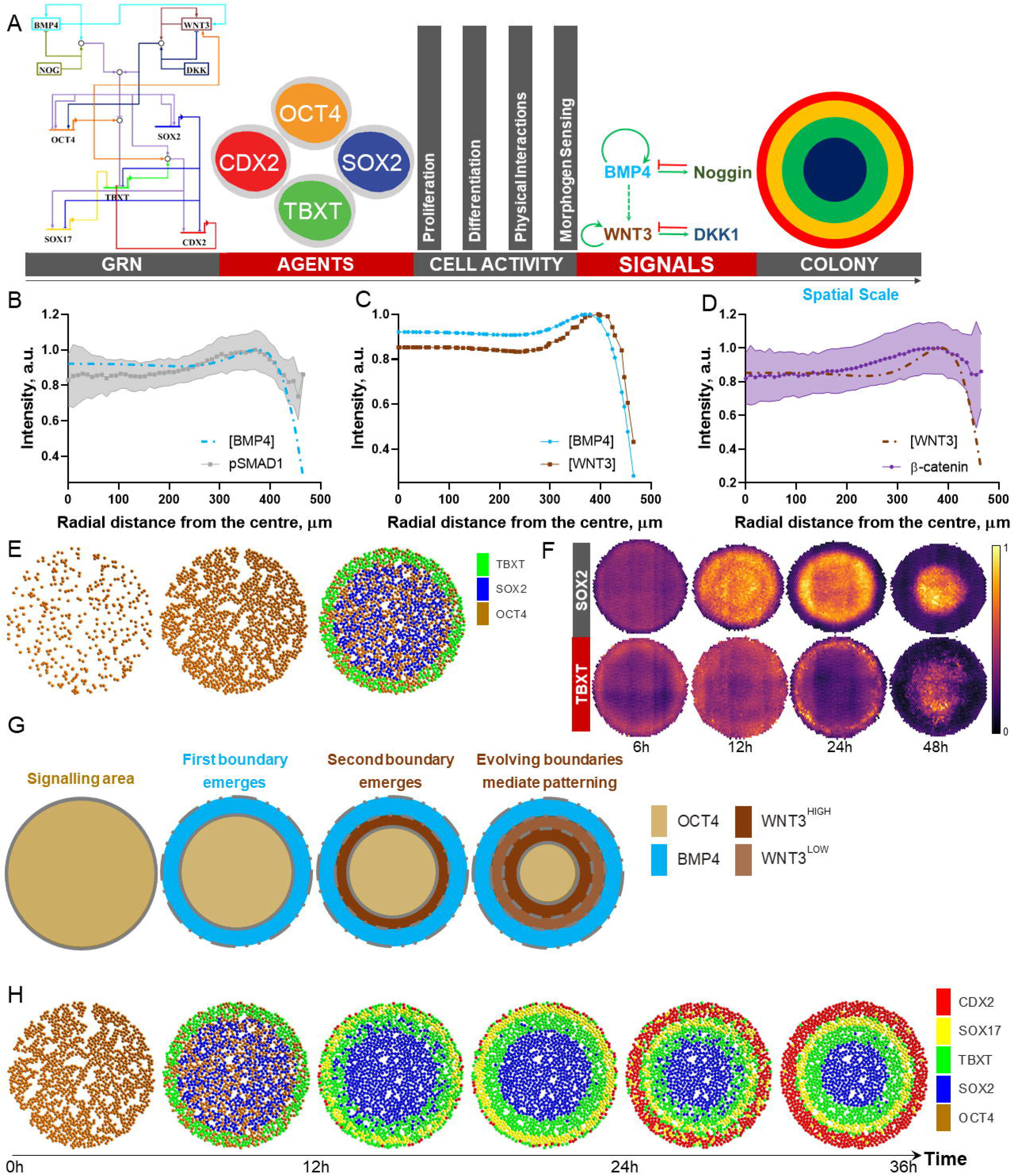
The virtual cell in a virtual environment paradigm. A. Schematic representing GARMEN’s architecture and the various decision-making hierarchies. B. Computational normalised [BMP4] vs empirical normalised pSMAD1 nuclear levels (n=29). Empirical data in solid (with standard deviation as error band), computational in dashed line. C. Computational normalised [WNT3] vs [BMP4]. D. Computational normalised [WNT3] vs empirical normalised β-catenin nuclear levels (n=29). Empirical data in solid (with standard deviation as error band), computational in dashed line. E. Simulation output capturing the first patterning phase: proliferation of cells until colony achieves confluence followed by the emergence of TBXT+ cells at the edge. F. Heatmaps showing the time evolution of mean SOX2 and TBXT expression in hPSC colonies treated with BMP4+NODAL at different time points: 6 h (n=29), 12 h (n=13), 24 h (n=15), and 48 h (n=27). The figure captures the radial internalisation of SOX2 and TBXT markers over time. G. Visualising the *step-wise mechanism* of radial internalisation of germ-layer markers. This mechanism mediates the internalisation of signaling boundaries in micropatterned hPSC colonies. H. Simulation output capturing the second and third peri-gastrulation patterning phases: radial internalisation of SOX2 & TBXT marker expression and the emergence of SOX17 ring, followed by the radial internalisation of SOX2, TBXT & SOX17 marker expression and the emergence of the CDX2 expression ring. n = number of replicates. Each counted replicate represents a circular hPSC colony.

**BOX 1:**
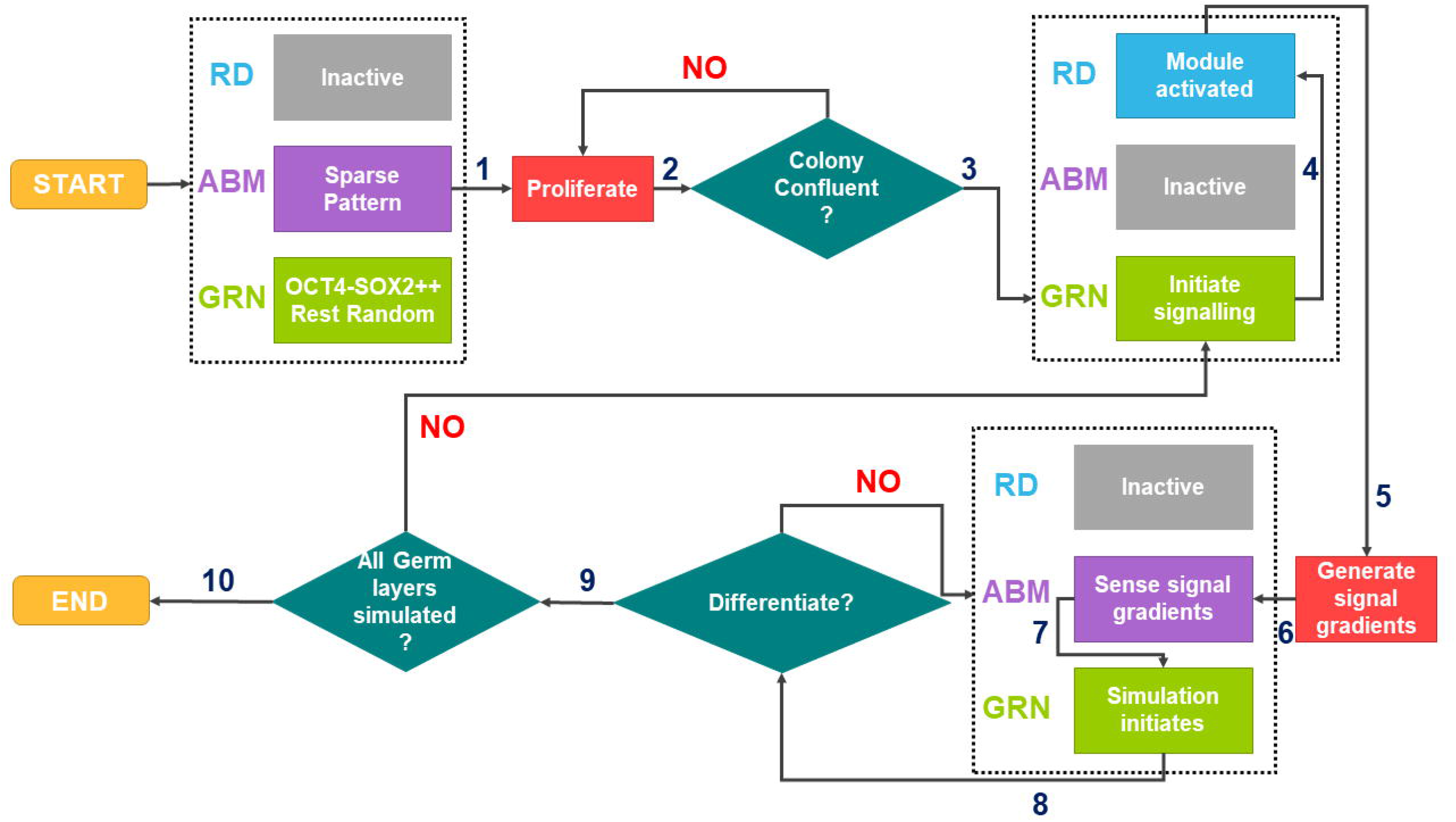
The GARMEN decision-making flowchart.

The arrows represent the sequence of decision-making steps in GARMEN. At the beginning of the simulation in GARMEN, few pluripotent agents populate the virtual colony. Simulation is initiated by the proliferation of these agents (Step 1). Confluence is automatically detected by the code when all agents have 4 or more neighbors. At this point (Step 2), the GRN is activated by triggering the BMP pathway (Step 3), which mimics the addition of BMP4 to the culture well. This activates the RD module, which calculates the concentration gradients (Step 4). Once the RD module reaches steady state, the spatial concentration profile (absolute values) of the signals is tabulated (Step 5) and passed to the agents (Step 6) that further bin it into ‘HIGH’, ‘LOW’, ‘OFF’ categories and pass it to the GRN (Step 7). The GRN is subsequently simulated so it can lead to relevant attractor states. The output from the GRN is fed back to agents at each iteration of the agent-based model, which, based on the states of each node and a differentiation probability, decide their phenotype per rules shown in **Methods** and **Table S4** (Step 8). Once agent ‘differentiation’ has reached steady state (~30 iterations, equivalent to 12 h in real time), we check if agents that are non-mesendodermal (or with CDX2 = OFF) have OCT4 either set to HIGH or LOW *and* whether BMP4 is present in the system, which is needed for the translocation of signals (Step 9). If these criteria are met, we assume that all possible germ-layers have *not* been observed. This leads to recomputing of the RD model with given new agent coordinates (Step 4), and so on. Finally, the simulation terminates when all possible germ-layers have been simulated (Step 10). A key point to note is that the transmission of information between the agent/GRN and RD layers (i.e. Steps 4, 5, and 6) as well as Step 9 are conducted manually in the current version of GARMEN. Lastly, as per the specific questions addressed in this work, our simulations only capture outcome up to 36 h post BMP4 stimulation.

We next simulated the model in the multi-paradigm *GARMEN* with 1200 virtual PSCs randomly located on a virtual circular micropattern. As initial conditions, we switched all the GRN nodes OFF, except OCT4 and SOX2, which were set to HIGH. BMP and WNT pathways were kept OFF to mimic experimental conditions (Tewary *et al*., 2017; Tewary *et al*., 2019). The virtual cells first initiated proliferation until the colony became confluent, achieved when each agent has at least 4 neighbours (see **Methods**). Following confluence, we mimicked the experimental induction of BMP4 by triggering the RD module. The concentrations of BMP4, NOGGIN and subsequently other morphogens, derived from the RD model were then relayed to agents, which based on their location inside the virtual colony received signal that was passed to the GRN (see **Table 1**).

To test whether the RD signaling pathway activation simulated was biologically relevant, we used a two-step approach. First, we ensured simulated radial profiles of morphogen activity (BMP4 and WNT3) were consistent with in vitro profiles of their transcriptional effectors (pSMAD1 and β-catenin, respectively). To achieve this, we formulated a quantitative metric wherein the radial locus of peak morphogen concentration (determined computationally) was compared to the mean radial locus of mean peak marker expression (**peak locus**, henceforth) determined experimentally. Comparing the computational (BMP4/WNT3) and experimental (pSMAD1/β-catenin) radial profiles served as an index of qualitative accuracy. The computationally predicted [BMP4] and [WNT3] profiles and peaks matched those of their empirical transcriptional effectors pSMAD1 and β-catenin (**Figures 2B-2D**) in vitro, respectively. Second, we tested whether the key parameter that led to these computational profiles were biologically relevant. We achieved this by comparing the effective diffusivity ratios of bmp4:noggin and wnt3:dkk used in the RD model (0.4 and 0.36, respectively) with their experimental estimates (0.38-0.77 and 0.34-0.36, respectively, detailed in **Methods**).

When these signal profiles were fed back to the GRN via the agents, the nuclear decision-making resulted in the emergence of TBXT agents at the colony edge where [BMP4] > [NOGGIN] (**Figure 2E**), thus capturing first of the three patterning phases. At this point, the RD module had reached its steady state, and so the signal gradients and, by extension, germ-layer patterns could no longer evolve. To drive the RD module out of its steady state and capture the radial internalisation of germ-layer markers, we focused on identifying the appropriate perturbation. Given the temporal emergence of patterns in these colonies reflects a changing boundary whereby TBXT expression emerges at the edge and then radially internalises (**Figure 2F**) (with SOX17 and CDX2 expression trailing this internalisation), we focused on altering the boundary of the geometry where the signalling module will remain active.

Given that TBXT expression is regulated when OCT4 interacts with high levels of BMP4/WNT3 (Funa *et al*., 2015; Wang *et al*., 2012), we reasoned that the radial internalisation of TBXT expression would result from shifting BMP4 or WNT3 activity. We hypothesised that at the spatial locus where [BMP4] and [WNT3] peak, OCT4 expression would be inhibited due to the emergence of TBXT+ cells. This creates a ‘boundary’ partitioning the space into OCT4^high^/undifferentiated (radially inwards of the boundary) and OCT4^low^/differentiated regions (radially outwards). Further, given WNT’s role in mediating OCT4 expression (*Sato et al., 2004*), we rationalised that the OCT4^high^ cells will remain engaged in WNT activity, which will in turn yield a new [WNT3] peak, creating a new boundary further separating OCT4^high^ and OCT4^low^ cells. At the systems-level, this constitutes a progressive OCT4^high^ expression front that translocates radially inwards. Consequently, this *step-wise mechanism* of internalisation of OCT4 expression, visualised in **Figure 2G**, due to mutually dynamic interplay between the gene network and morphogen signals leads to radial internalisation of germ-layer patterns.

We next applied this *step-wise* mechanism to our GARMEN model to capture the remaining patterning phases. The mechanism was implemented by identifying the boundary separating undifferentiated vs differentiated virtual cells from the agent-based model, using it to re-solve the RD model to its steady state (with the new boundary), and relaying the new gradients to the GRNs via agents. The new gradients were obtained by simulating the RD equations fully for the region with undifferentiated agents but setting certain WNT module reaction kinetics to zero for the regions with differentiated agents (see **Methods**). In the agent-based model, continuing from the first phase, the internal non TBXT agents within the radial distance of 380 μm continued to engage in full WNT reaction-diffusion. When the agents were fed with the new [morphogen] profiles we observed the loss of SOX2 agents close to the initial boundary, giving way to TBXT agents. SOX2 agents, thus, internalised radially, and TBXT agents along with it (**Figure 2H**, **Movie S1**). We also noticed the emergence of SOX17 agents at the colony edge, due to them displaying a low OCT4 state in combination with high [BMP4]. This observation was consistent with empirical observations and reminiscent of the second patterning phase (**Figure 2H**). The model once again reached a steady state, giving us the new boundary at 300 μm. The RD module was simulated again with this new boundary (with only the undifferentiated TBXT–SOX17–agents continuing full WNT activity). This led to a further internalisation of the germ-layer states and the emergence of a CDX2 agent population at the edge (**Figure 2H**, **Movie S1**). This marked the third and final patterning phase, thereby capturing not only the precise spatial but also temporal order of germ-layer states as well as the radial internalisation of signalling activity.

Taken together these results support our model’s ability to capture emergence of spatial patterns in both space and time based on the activity of individual GRNs. These results also suggest that crosstalk between signalling pathways (specifically BMP and WNT) contextualises the differentiation signalling cues for gene networks during germ layer emergence. This contextualising effect is best evidenced by the lack of peri-gastrulation patterns when GARMEN is run without the RD module (**Figure S3A**). We next tested the predictive capability of our framework by exploring the peri-gastrulation pattern phase space.

### Exploring the peri-gastrulation pattern phase space

The predictive capability of GARMEN was tested by simulating germ-layer patterns due to (single and multiple) perturbations in the GRN. This entailed setting the activity levels of the relevant nodes continuously to OFF (represented as ‘–’ in **Figure 3A** reflecting in vitro gene knockout) or HIGH (represented as ‘++’ reflecting in vitro ectopic expression). Single perturbations involved changing activity levels of a single node (e.g. OCT4−, etc.), whereas multiple perturbations involved changing activity levels of multiple nodes (i.e. OCT4− & SOX2++ & TBXT−, etc.). The pattern phase space is represented in **Figure 3A**.

**Figure 3.**
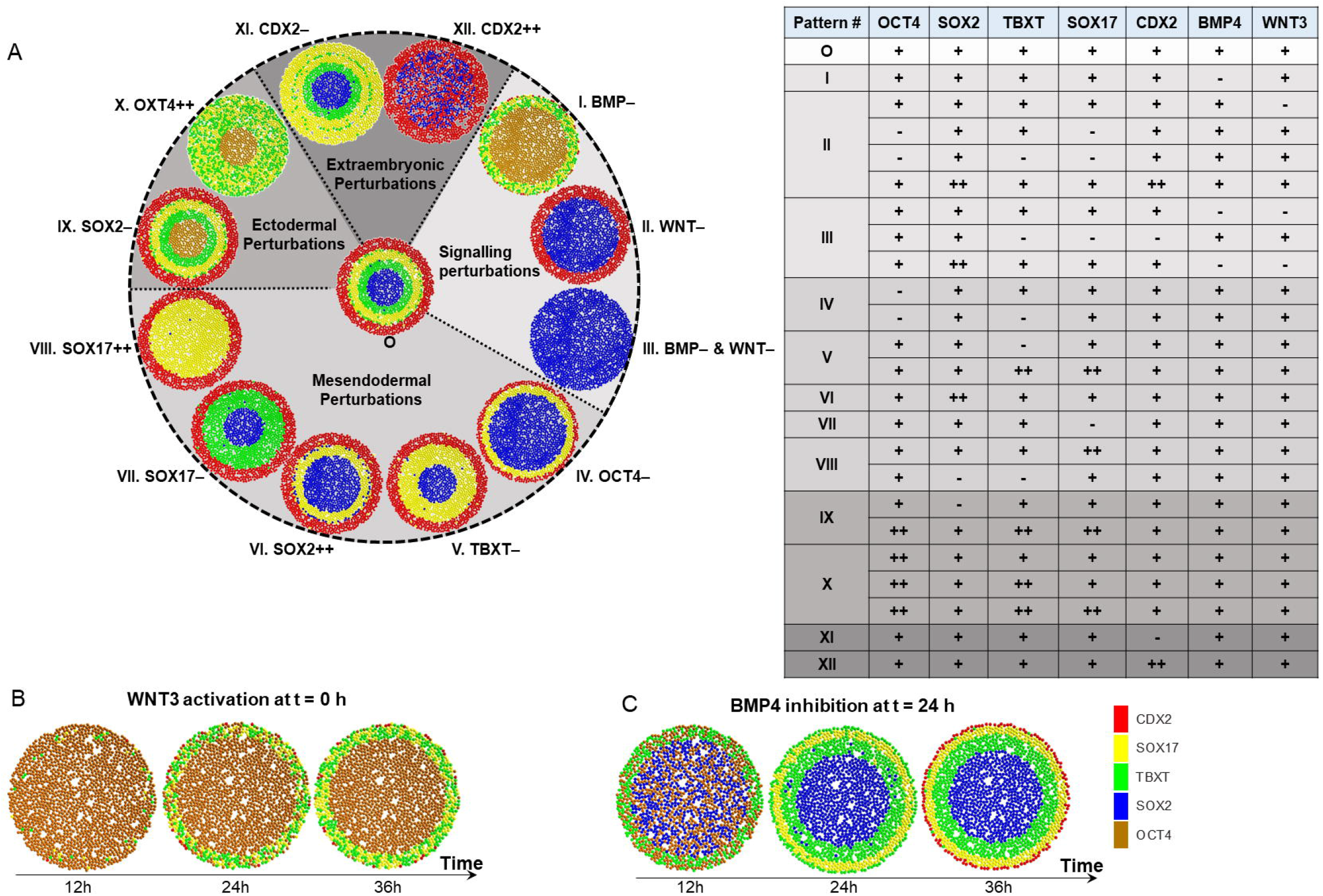
Exploring the peri-gastrulation phase space. A. Results of the parametric analysis conducted to link network structure to the germ-layer pattern phase-space. The table represents the perturbation matrix that led to the various patterns. ‘–’ indicates the node was continuously set to OFF, ‘++’ means node was continuously set to HIGH, whereas ‘+’ means node could vary between OFF, LOW, or HIGH. B. Simulation output when the model bypasses BMP activity by initiating with WNT activity. C. Simulation output when BMP activity is inhibited midway (equal to 24 h of real time) during patterning.

**Figure 3A** also shows the impact of perturbations on the properties of the patterns (i.e. how many germ layers are observed and ratio of various germ layers within a pattern). We noted that in terms of signalling the loss of WNT pathway had the most severe impact on patterning. Specifically, BMP4− led to the emergence of all germ-layer states albeit with a lower number of mesendodermal agents (790, vs 621 agents in Pattern O). However, WNT3− led to the loss of germ-layer pattern yielding only a SOX2 expressing colony core and CDX2 expressing edge (Pattern II), consistent with earlier observations (Martyn *et al*., 2018). WNT3− and BMP4− together led to a total loss of annular patterns (Pattern III), which is also consistent with earlier reports that lack of exogenous signals results in the acquisition of ectodermal fate (Fathi, Eisa-Beygi and Baharvand, 2017; Blauwkamp *et al*., 2012; Smukler *et al*., 2006). We also tested an ‘inverse’ condition whereby a strong BMP4 signal was imposed at the centre of the colony as opposed to the edge. This led to the emergence of the mesendoderm at the centre of the colony (**Figure S3B**), which has been observed in vitro under similar conditions (Regier *et al*., 2019).

We next perturbed the pluripotency and mesendodermal nodes (similar patterns are reported together). OCT4−perturbation with BMP4+ led to the removal of TBXT expressing agents, but preserved the SOX2, SOX17, CDX2 displaying radial agent order (Pattern IV). This overall pattern is also observed with TBXT– (Pattern V); OCT4−/TBXT– (Pattern IV); and SOX2++ (Pattern VI) perturbations. SOX17− led to the emergence of ectodermal−, mesodermal−, and extraembryonic-associated states (Pattern VII). SOX17++ on the other hand only yielded endodermal and extraembryonic states (Pattern VIII). SOX2− led to the emergence of mesendodermal and extaembryonic states, but not ectodermal states (Pattern IX). OCT4++, on the other hand, suppressed the ectodermal-associated phenotype but permitted mesendodermal-associated phenotype (Pattern X). TBXT++ resulted in the same patterns as the original condition (Pattern O), though it may potentially limit the number of SOX2 and SOX17 agents in the pattern. Further, CDX2− resulted in the three germ-layer rings without the CDX2 ring at the edge (Pattern XI), whereas CDX2++ only yielded ectodermal and extraembryonic cell states (Pattern XII). We anticipate the SOX2 expressing virtual cells here represent extraembryonic ectodermal state.

It is noteworthy that with our minimal GRN and its logic (**Table 1**) the pattern phase space is finite and repetitive. No new patterns emerged with added combinations of simulated knockdowns or ectopic expressions of multiple GRN components. For example, TBXT– & SOX17– or SOX17– & OCT4− yield Pattern II; CDX2− & TBXT− leads to Pattern VIII; CDX2− & TBXT− & SOX17− will result in Pattern III. Additionally, TBXT++ & OCT4++ will yield Pattern X, whereas SOX17++ & TBXT++ will lead to Pattern V, and so on. All perturbations and the resulting patterns are tabulated in **Figure 3A**. Despite a potentially vast perturbation combinatorial space, the number of eventual germ-layer patterns is finite due to the limited context offered by these combinations and constraints outlined as initial conditions (in **Figure 3A**) and GRN rules (in **Table 1**). This effectively constitutes GRN pruning that limits the number of single-agent states, which collectively form a limited number of patterns when agents receive input from the RD module.

We further observed in the BMP4– simulation that the TBXT state was confined to the colony edge, which did not internalise radially for the duration of the simulation (**Figure 3B**, **Movie S2**). In fact, the simulation showed SOX17 and TBXT agents that were co-localised, possibly suggesting this condition does not commit mesendodermal agents to either mesodermal or endodermal states. Curiously, we also note a wide spread of agents expressing OCT4 at the colony centre and reasoned this to be a result of absence of BMP activity and the presence of WNT activity. This suggested that the peri-gastrulation patterns evolve as a result of the interplay between OCT4 and BMP & WNT pathways (as encapsulated by the proposed *step-wise* mechanism).

To further investigate this interplay, we tested the consequence of halting the translocating differentiation front. We initiated the model by activating the BMP module, but turning the BMP activity OFF after 24 hours (h) in the simulation (**Figure 3C**). In this simulation, while we noticed a shift in TBXT state radially inwards during the 1^st^ and 2^nd^ pattering phases (i.e. 12-24 h), there was no shift after the BMP module was switched off (**Figure 3C**, **Movie S3**), suggesting that peak locus of TBXT state at 48 h should not be different than after 24 h of exposure to BMP4. We also observed SOX17- and CDX2-asociated agents that were both confined to the colony edge.

In summary, we used a validated hPSC lineage commitment GRN structure to explore relationship between GRN wiring and the peri-gastrulation pattern phase space. We also found that an inward travelling differentiation front can be predictively interrupted by altering the input signals to the virtual colonies. The simulations also revealed that BMP4 is key to initiating this front, which stays static in the absence of BMP4. The simulations predicted that bypassing BMP activity limits mesendoderm phenotype to colony edge, and inhibiting BMP activity mid-way during experimentation halts the translocating front entirely. These predictions were subsequently tested in vitro.

### Lineage commitment during peri-gastrulation patterning is mediated by dynamic signalling boundaries regulated by OCT4, WNT3 and BMP4 interplay

We next set out to examine whether the OCT4, BMP4, and WNT3 interplay as observed in the model and its impact on germ layer-like patterning was reflected in the hPSC colonies. We set out to test this by first predicting the empirical profile of OCT4 based on the radial [BMP4] profile at 12 h. Per our model, to yield TBXT expression at the observed radial locus given [BMP4] (**Figure 4A**), OCT4 expression should be low at the colony core, increase with increasing [BMP4], and then drop towards the edge (**Figure 4A**). SOX2 on the other hand will be high at the core (where [BMP4] < [NOGGIN]) but fall in the region where [BMP4] peaks. This was confirmed experimentally (**Figure 4B**).

**Figure 4.**
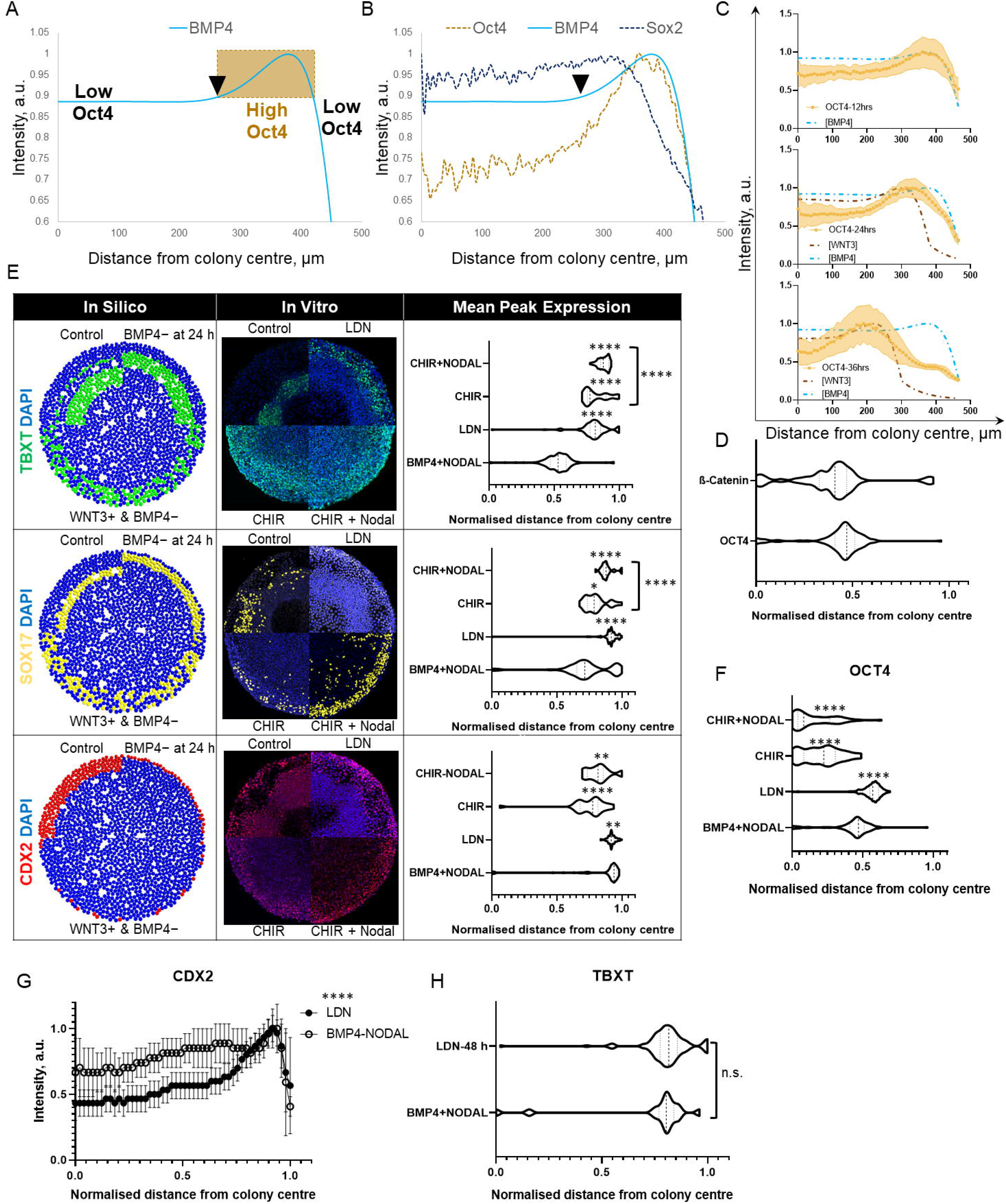
WNT activity with OCT4 expression directly mediates radial internalisation in the presence of BMP4. A. Computational radial [BMP4] profile (solid line). The arrowhead marks the spot where [BMP4] profile begins to increase in intensity, which we predicted will lead to increase in OCT4 expression profile. The fall in [BMP4] following the peak was predicted to again correlate with LOW OCT4 expression. Both predictions were based visually on the [BMP4] profile. B. Comparison of in vitro (OCT4 and SOX2) and computational (BMP4) data to test the prediction indicated in A. Dotted lines represent empirical data, solid lines represent computational data. C. Comparison of OCT4 peak loci at 12 h (n=18), 24 h (n=19), and 36 h (n=33) against computational [BMP4] and [WNT3] profiles. Solid lines with square show mean OCT4 expression with error band representing standard deviation. Dash-dotted lines show computational data. D. Comparison of the peak loci of β-catenin (n=151) and OCT4 (n=118) in the ‘control’ condition after 48 h. E. Comparison of in silico (left column) predictions against in vitro data (mid and right columns). The images under the mid column represent a composite of hPSC colonies maintained under different conditions. The plots under the right column compare the peak loci of indicated markers for the four conditions. p-values represent comparison against the control condition, except where direct comparison between CHIR and CHIR+NODAL are specified. For TBXT: n=118 (control), 127 (LDN), 35 (CHIR), and 63 (CHIR+NODAL). For SOX17: n=75 (control), 73 (LDN), 28 (CHIR), and 19 (CHIR+NODAL). For CDX2: n=75 (control), 73 (LDN), 28 (CHIR), and 19 (CHIR+NODAL). F. Comparison of OCT4 peak loci for the four conditions in E. p-values represent comparison against the control condition. n=118 (control), 127 (LDN), 35 (CHIR), and 63 (CHIR+NODAL). G. Radial CDX2 expression profile (mean values represented by circles and standard deviation by error bars) comparison between the control (n=75) vs LDN (n=63) treated hPSC colonies. H. Comparison of the peak loci of TBXT expression in control at 24 h (n=16) vs LDN (n=127) treated cells after 48 h. p = 0.67. * p<0.05; ** p<0.01; *** p<0.001; **** p<0.0001; ns, not significant; n, number of colonies. Refer to **Methods** for details of statistical tests conducted.

Beyond this snapshot, we next tested whether we can predict OCT4 expression profile over time based on our computational [BMP4] and [WNT3] profiles. We achieved this by comparing OCT4 peak locus with radial loci of peak [BMP4] and [WNT3] across the three patterning phases. The [BMP4] and [WNT3] profiles with the corresponding OCT4 expression over 36 h is shown in **Figure 4C**. The figure shows close agreement between the (qualitative) shape of the computational & empirical profiles (i.e. profiles show ascent and descent at similar radial loci) and (quantitative) peak loci of OCT4 expression and [BMP4] & [WNT3] across 36 h. To further confirm this interplay, we compared (**Figure 4D**) the radial profiles of β-catenin vs OCT4 at the end of the experiment (i.e. 48 h post differentiation). We observed these to show a similar peak location and radial profile, with β-catenin peaking just inside of OCT4 (mean ± standard deviation peak loci = 190±95 μm vs 210±70 μm, respectively).

We next tested whether based on the model we can predictively disrupt the internalisation of the mesendodermal markers in vitro. Our model had predicted that bypassing BMP activity will limit the mesendoderm-associated (TBXT/SOX17) agents towards the edge vs when BMP4 was used to trigger differentiation (**Figure 4E**). To test this prediction, we initiated differentiation in vitro by triggering the WNT pathway directly via the WNT agonist CHIR-99021 (CHIR). hPSC colonies exposed to both CHIR and CHIR+NODAL showed the TBXT & SOX17 peak locus significantly closer to the edge (**Figure 4E**, all p-values indicated in the figure henceforth) compared to the colonies exposed to BMP4+NODAL (control). We noted that the two conditions led to different spatial extent of SOX17 expression: CHIR led to a thin spread of SOX17+ cells whereas CHIR+NODAL resulted in a wider spread. We also observed that the mean peak TBXT and SOX17 loci for CHIR+NODAL was significantly closer to the edge vs CHIR. This is consistent with our proposed *step-wise mechanism* that due to NODAL’s BMP inhibitory effect (Tewary *et al*., 2019) the TBXT and SOX17 peak locus will be confined closer to the edge compared to a condition where BMP inhibitor is not used (CHIR in this case).

Our simulations also showed that bypassing BMP activity leads to co-localization of TBXT and SOX17 agents (**Figure 3B**), indicating that divergence to mesoderm and endoderm had not occurred. This was explored experimentally by comparing the peak loci of TBXT vs SOX17 for the two CHIR conditions (**Figures S4A and S4B**) that were not significantly different. This was in stark contrast against the control condition where the mean peak locus of TBXT was significantly closer to the colony centre vs SOX17, suggesting that the control condition results in the divergence between mesoderm- and endoderm-associated markers (**Figure S4C**). Importantly, colonies stimulated with CHIR and CHIR+NODAL showed a wider spread of OCT4 expressing cells vs control (**Figures 4F, S4D, S4E**), as suggested in silico (**Figure 3B**).

Next, we tested the impact of suppressing BMP activity mid-way through the experiment, which our model predicted will limit the mean peak expression of the mesendodermal markers to the time of exposure to inhibitor (**Figure 3C**). This was tested by initially stimulating the hPSC colonies with BMP4 and NODAL but switching to media containing the BMP inhibitor LDN193189 (LDN) for 24 h. The LDN containing media was devoid of BMP4 and NODAL. Consistent with our prediction, we noticed the peak expression of TBXT confined significantly closer to the edge vs control (**Figure 4E and S4F**). Upon measuring the peak SOX17 expression we noticed it was also significantly confined to the edge vs control (**Figure 4E and S4G**).

Finally, we compared the peak locus of CDX2. The model predicted that the control condition will have CDX2-associated agents limited to the edge, albeit with a larger spatial spread that extends to colony centre (**Figure 4E**). It also predicted that inhibiting the BMP pathway midway will lead to CDX2-associated agents confined to the edge (**Figure 4E**). In vitro, we found that the peak CDX2 locus for both the control and LDN conditions was located close to the edge (**Figure 4E**). However, the two conditions yielded significantly different data. To understand this, we examined the radial CDX2 expression profiles for both conditions (**Figure 4G**) and noted that CDX2 expression remained significantly high towards the colony centre for control hPSC colonies vs the LDN treatment. This was consistent with our model (**Figure 4E**).

Finally, to confirm that we were truly able to halt the internalisation, we compared the radial locus of peak TBXT expression of hPSC colonies kept in the control media for 24 h vs those exposed to LDN for 24 h (after 24 h of BMP4 exposure). As predicted by the model that the TBXT state will not translocate internally (between 24 h and 36 h in **Figure 3C**), the peak TBXT loci for these two cases, shown in **Figure 4H**, were not significantly different. Finally, like control, the spatial correlation between the radial OCT4 and β-catenin expression profiles was also observed for this condition (**Figure S4H**).

These observations, especially our ability to predict the position of the travelling OCT4 expression front based on computational [BMP4] & [WNT3] profiles, suggests a possible interplay between OCT4 and BMP & WNT pathways and supports the *step-wise mechanism* of translocating OCT4 expression front. Our results further show that our minimalist GRN-based multi-paradigm model can capture spatial germ-layer patterning based on gene activity at individual cell-level with high fidelity.

## DISCUSSION

In this study, we develop a multi-paradigm platform, GARMEN, to investigate the mechanisms mediating the emergence of peri-gastrulation patterning in geometrically confined hPSC colonies. The platform integrates three hierarchical levels of cell fate control: regulatory, cellular, and microenvironmental. We achieved this by embedding a minimalist PSC GRN into ‘cellular’ agents that were contextualised in a biochemically dynamic microenvironment. GARMEN allowed us to capture both the spatiotemporal signalling dynamics and germ-layer patterns with high fidelity as a function of the GRN structure within individual cells interacting via paracrine signalling activity. Using GARMEN, we quantitatively reveal that mutually reciprocal dynamic crosstalk between the GRNs and multiple signal pathways regulate germlayer patterning. Specifically, the single-cell GRNs collectively shape the global and local signalling profiles, which in turn engage the single-cell GRNs back by providing the local context to undergo differentiation, which in turn alters the signalling profile, and so on.

Specific to germ layer emergence in micropatterned hPSCs, our work reveals how the environmental BMP4 and WNT3 signals are processed by the GRN, in each cell in a spatially differentiated manner to yield germ-layer patterns (**Figure 5A**). In this regard we show how sequential BMP4-OCT4 and, subsequently, WNT3− OCT4 interactions result in an inwardly translocating OCT4 expression front due to the formation of boundaries that separate the colony into OCT4^high^ and OCT4^low^ regions. This front contributes to a continuously changing WNT signalling profile, evidenced by the radial internalisation of β-catenin in hPSC colonies (**Figure 5B**). While dynamically altering biological boundary conditions are conceptually (Vahey and Fletcher, 2014) accepted to play a key role in guiding development, they have rarely (Chen *et al*., 2018) been observed in silico and validated in vitro. Further, Cotterell *et al*. (2015) had previously explained the existence of a progressive differentiation front in somites via a networkbased reaction-diffusion model to create a periodic pattern of expression stripes. Our approach advances the complexity and application of this paradigm using agents. We also capture how crosstalk between the four signal molecules contextualise differential cues spatially. For example, in our model the spatial region where WNT3>DKK and BMP4<NOGGIN drive OCT4 state, whereas the ectoderm state dominates in the region where both WNT3 & BMP4<DKK & NOGGIN.

**Figure 5.**
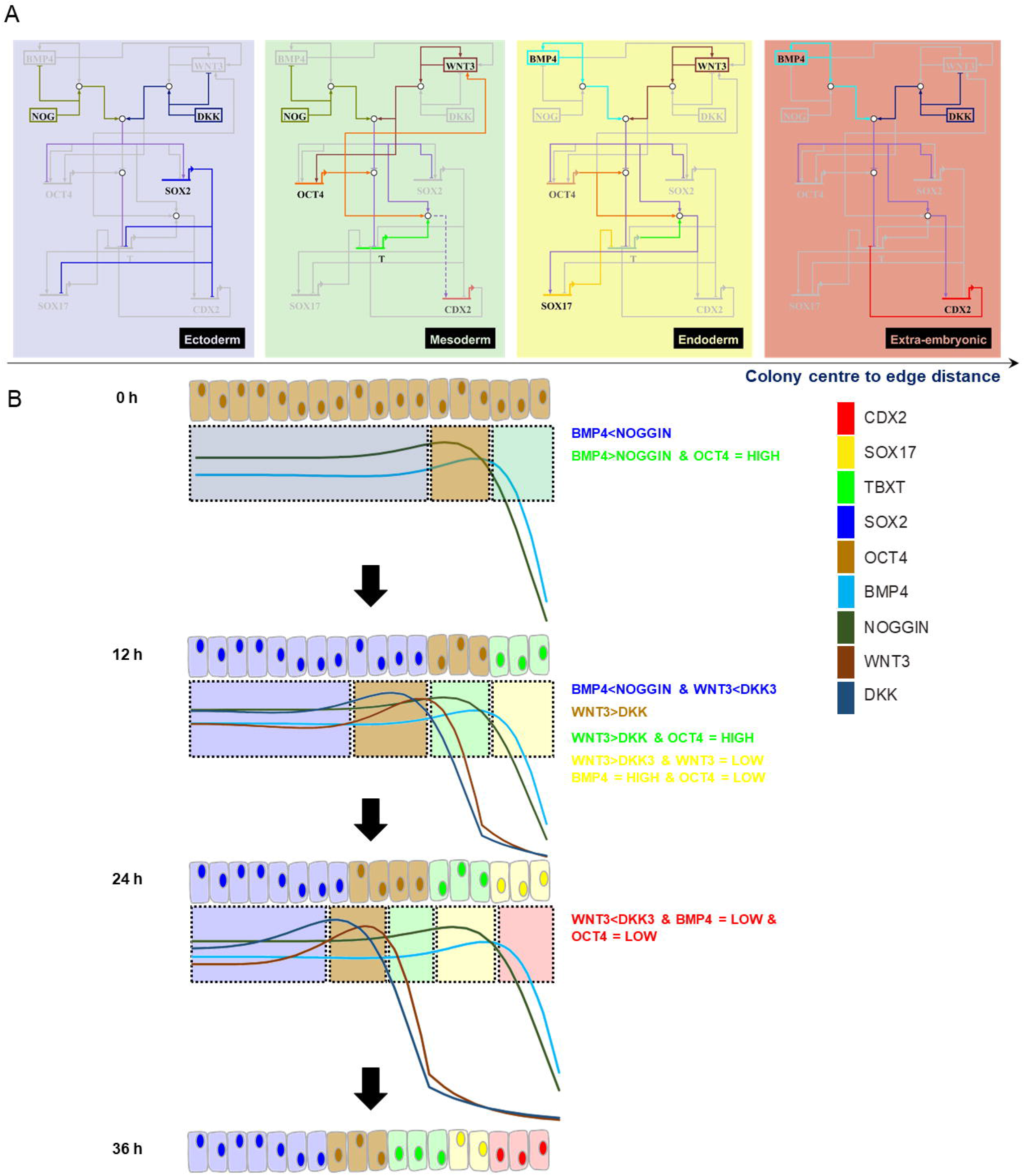
Peri-gastrulation patterning is mediated by spatial signals that engage the GRN in a spatially differential manner. A. The figure captures the contextualising effect of exogenous signals on single-cell GRNs as the hPSC colony undergoes peri-gastrulation patterning. Different germ-layer fates are symbolised with coloured boxes. Active genes and interactions that lead to a particular germ-layer fate are shown in colour, whereas genes and interactions that are inactivated due to the signalling context are shown in gray. This schematic highlights how different signal inputs activate and silence different topological paths within an unchanging GRN to reveal the spatially distinct peri-gastrulation patterns. B. The figure summarises how the single-cell GRNs collectively influence their signalling profiles over time, which in turn contextualise the GRN to yield germ-layer patterns.

Additionally, our model captures the space and time sequence of peri-gastrulation pattern phases. This is supported by our exploration of the germ-layer pattern phase space, which predicted the perturbations that will modify the patterns (by bypassing BMP pathway) and making the system abrogate patterning (by suppressing the BMP pathway). Consequently, our model explains the majority of observations made to date using the 2D hPSC colonies (as reported in *Results*). It additionally explains why, in case a part of the colony (e.g. the core) is continuously exposed to continuous BMP4 or WNT3 profile, it will express mesendodermal markers compared to its neighbours with relatively low BMP4/WNT3 exposure (Regier *et al*., 2019; Manfrin *et al*., 2019). Critically, our results build on a recently proposed biophysical mechanism of initiating TBXT expression in micropatterned hPSC colonies (Muncie *et al*., 2020) and explain how it links with the reaction-diffusion paradigm to yield peri-gastrulation patterns. Specifically, Muncie *et al*. (2020) propose that high cell-adhesion tension promotes the release of β-catenin from cadherin-catenin complexes, which enhances mesoderm-inducing WNT signalling. Our study suggests that this subsequently emerging WNT activity leads to the second and third patterning phases (as explained in **Figure 2G**) in the hPSC colonies. Our work, thus, offers a glimpse into how the biochemical and mechanical cues together regulate peri-gastrulation patterning.

Models that have previously tried to explain the emergence of germ-layer patterns have relied on gradients of individual signals including BMP4 (Etoc *et al*., 2016; Tewary *et al*., 2017), NODAL (Tewary *et al*., 2019), and WNT (Chhabra *et al*., 2019). We build on these models in three ways. First, we modelled an integration of multiple pathways that considered the concentrations of multiple activator (BMP4 and WNT3) and inhibitor (NOGGIN and DKK) signals. Second, we simulated the spatiotemporal evolution of germ-layer patterns as a consequence of dynamic interactions and activities of the underlying decision-making components (agents) that make up the simulated tissue. This is why the GRN by itself and GRN within agents but without the RD module (**Figure S3A**) cannot capture spatial and temporal patterning aspects at the level of germ-layer markers. Third, the patterns as they arise are *dynamically relevant* (Rosas *et al*., 2020), i.e. they contain the information that predicts future evolution of the system. For example, we need to know the status of WNT signal active agents (at 12 and/or 24 h, for example) to predict the temporal evolution of peri-gastrulation patterning at the subsequent time steps (**Figures 2H, 4C, and 4D**). Together these aspects start to capture elements of multicellular emergent systems. Embedding the morphogen gradient computing equations in each agent to allow for biochemical feedback between individual agents, as has been done elsewhere (Kaul, Cui and Ventikos, 2013; Kaul and Ventikos, 2015a), will expand this capability further. This is significant as our approach can be coupled with GRNs developed using (e.g. bulk−, single cell−, ATAC−) sequencing data to quantitatively understand two fundamental questions: how do biological function (i.e. *how does collective cell behaviour yield organised tissues?*) and pathology (i.e. *how changes in gene activity mediate disease phenotypes?) emerge* at the tissue/organ levels. This will allow us to engineer strategies that will enable development of optimal regenerative therapies (Tewary, Shakiba and Zandstra, 2018; Prochazka, Benenson and Zandstra, 2017) and conduct patient-specific modelling to help suggest interventions tailored to the patients’ genomic profile (Kaul, 2019).

While the model can predict patterning for different initial and boundary conditions, it has limitations. First, the model does not consider the impact of mechanical forces during selforganisation (Muncie *et al*., 2020; Xue *et al*., 2018). Mechanical forces are potentially responsible for the localised increase in height in hPSC colonies observed around the annulus where cells are expressing TBXT. Therefore, the model does not accurately predict events beyond 36 h when the colony depth reaches several cell layers. Second, the model can currently only capture peri-gastrulation patterns in 2D. This is again due to lack of a mechanical module in this platform. Third, our model did not capture the spatial differences between the CHIR vs CHIR+NODAL conditions (with peak marker loci for CHIR+NODAL significantly different from CHIR stimulation). This discrepancy can be explained by our assumption in the model that BMP activity could not be switched on downstream if it is initially bypassed, which is similar to the CHIR+NODAL effect where NODAL acts as a BMP inhibitor. Despite this, our model still captured the exteriorising effect of bypassing (via CHIR) and inhibiting (via NODAL and LDN) BMP activity on the patterns. Finally, this initial iteration of our platform did not consider NODAL and its interactions with the various genes represented in our GRN, but the modularity of our approach means this can be easily added.

Our approach does, however, reveal how a limited number of signalling pathways in a confined space with a developmentally homogeneous cell population can yield a complex structure. While we employed a highly abstracted GRN, it serves as a starting point to understand how gene activity mediates spatial patterns in cell populations and it has the potential to reveal how the various signalling pathways are repurposed as the embryo grows in size to reveal multiple niches needed for the formation of disparate tissues. Doing so entails broadening this GRN to capture more advanced post-gastrulation patterning events, by including additional signalling pathways (e.g. NODAL, NOTCH), genes (e.g. SNAIL, OTX2, RUNX1), and explicit epigenomic components (e.g. to account for DNA methylation, RNA silencing) as needed. As discussed above, our approach also offers a systematic way to integrate biochemical signalling with mechanical cues to understand post-gastrulation patterning. This further requires the ability to account for changes in size (in 3 dimensions), morphology, and migratory behaviour post biochemical stimulation. Doing so entails exploiting the *proliferation* and *physical interactions* modules in the framework and integrating them with a model of mechanical forces. This will enable GARMEN to calculate forces acting on each agent, due to each other or agent differentiation / proliferation, and consequently capture macroscopic phenomena such as cell migration, layer thickening, and geometric contraction or extension. This will also allow us to quantitatively understand emergent phenomena in its broadest context within developmental systems. We aim to address these aspects in the next iteration of our model, including adding machine learning algorithms that can analyse and self-direct the GRN towards organogenic patterns to explain known knowns and reveal unknown unknowns.

## ACKNOWLEDGEMENTS

We thank Dr Andras Nagy for the CA1 cell line. The project was supported by the CIHR Foundation Grant (FDN-154283). HK is currently supported by the Michael Smith Foundation for Health Research Trainee Award (Award #18427). NW is being supported by Alexander Graham Bell Canada Graduate Scholarship. AH is being supported by MITACS Elevate Award (Award # IT15865). We also acknowledge support from Dr Jennifer Ma towards editing of manuscript figures.

## AUTHOR CONTRIBUTIONS

Conceptualisation, H.K. and P.W.Z.; Methodology, H.K. and M.T.; Software, H.K.; Validation, H.K., J.O., and D.A.H; Formal Analysis, H.K., D.A.H, A.H., and J.O.; Investigation, H.K. and N.W.; Writing – Original Draft, H.K., N.W., A.H., and J.O.; Writing - Review & Editing, H.K., J.O., D.A.H., M.T., and P.W.Z.; Visualisation, H.K., J.O., and D.A.H.; Funding Acquisition, P.W.Z.

## REFERENCES

Abbott, R. G., Forrest, S. and Pienta, K. J. (2006). Simulating the hallmarks of cancer. Artif Life 12(4), 617–634.

Adra, S., Sun, T., MacNeil, S., Holcombe, M. and Smallwood, R. (2010). Development of a three dimensional multiscale computational model of the human epidermis. PLoS One 5(1), e8511.

An, G. (2004). In silico experiments of existing and hypothetical cytokine-directed clinical trials using agent-based modeling. Crit Care Med 32(10), 2050–2060.

Bailey, A. M., Thorne, B. C. and Peirce, S. M. (2007). Multi-cell agent-based simulation of the microvasculature to study the dynamics of circulating inflammatory cell trafficking. Ann Biomed Eng 35(6), 916–936.

Blauwkamp, T. A., Nigam, S., Ardehali, R., Weissman, I. L. and Nusse, R. (2012). Endogenous Wnt signalling in human embryonic stem cells generates an equilibrium of distinct lineage-specified progenitors. Nat Commun 3, 1070.

Boyer, L. A., Lee, T. I., Cole, M. F., Johnstone, S. E., Levine, S. S., Zucker, J. P., Guenther, M. G., Kumar, R. M., Murray, H. L., Jenner, R. G., Gifford, D. K., Melton, D. A., Jaenisch, R. and Young, R. A. (2005). Core transcriptional regulatory circuitry in human embryonic stem cells. Cell 122(6), 947–956.

Bérenguier, D., Chaouiya, C., Monteiro, P. T., Naldi, A., Remy, E., Thieffry, D. and Tichit, L. (2013). Dynamical modeling and analysis of large cellular regulatory networks. Chaos 23(2), 025114.

Casal, A., Sumen, C., Reddy, T. E., Alber, M. S. and Lee, P. P. (2005). Agent-based modeling of the context dependency in T cell recognition. J Theor Biol 236(4), 376–391.

Chachi, L., Diver, S., Kaul, H., Rebelatto, M. C., Boutrin, A., Nisa, P., Newbold, P. and Brightling, C. (2019). Computational modelling prediction and clinical validation of impact of benralizumab on airway smooth muscle mass in asthma. Eur Respir J 54(5), 1900930.

Chen, K., Vigliotti, A., Bacca, M., McMeeking, R. M., Deshpande, V. S. and Holmes, J. W. (2018). Role of boundary conditions in determining cell alignment in response to stretch. Proc Natl Acad Sci U S A 115(5), 986–991.

Chhabra, S., Liu, L., Goh, R., Kong, X. and Warmflash, A. (2019). Dissecting the dynamics of signaling events in the BMP, WNT, and NODAL cascade during self-organized fate patterning in human gastruloids. PLoS Biol 17(10), e3000498.

Coakley, S., Smallwood, R. and Holcombe, M. (2006). Using X-machines as a formal basis for describing agents in agent-based modelling. Spring Simulation Multiconference 2006, Huntsville, Alabama, USA, 33–40.

Collombet, S., van Oevelen, C., Sardina Ortega, J. L., Abou-Jaoudé, W., Di Stefano, B., Thomas-Chollier, M., Graf, T. and Thieffry, D. (2017). Logical modeling of lymphoid and myeloid cell specification and transdifferentiation. Proc Natl Acad Sci U S A 114(23), 5792–5799.

Cotterell, J., Robert-Moreno, A. and Sharpe, J. (2015). A local, self-organizing reaction-diffusion model can explain somite patterning in embryos. Cell Syst 1(4), 257–69.

Cotterell, J. and Sharpe, J. (2010). An atlas of gene regulatory networks reveals multiple three-gene mechanisms for interpreting morphogen gradients. Mol Syst Biol 6, 425.

Davidson, K. C., Adams, A. M., Goodson, J. M., McDonald, C. E., Potter, J. C., Berndt, J. D., Biechele, T. L., Taylor, R. J. and Moon, R. T. (2012). Wnt/β-catenin signaling promotes differentiation, not self-renewal, of human embryonic stem cells and is repressed by Oct4. Proc Natl Acad Sci U S A 109(12), 4485–4490.

Dunn, S. J., Li, M. A., Carbognin, E., Smith, A. and Martello, G. (2019). A common molecular logic determines embryonic stem cell self-renewal and reprogramming. EMBO J, 38(1), 100003.

Dunn, S. J., Martello, G., Yordanov, B., Emmott, S. and Smith, A. G. (2014). Defining an essential transcription factor program for naive pluripotency. Science 344(6188), 1156–1160.

Etoc, F., Metzger, J., Ruzo, A., Kirst, C., Yoney, A., Ozair, M. Z., Brivanlou, A. H. and Siggia, E. D. (2016). A balance between secreted inhibitors and edge sensing controls gastruloid selforganization. Dev Cell 39(3), 302–315.

Faial, T., Bernardo, A. S., Mendjan, S., Diamanti, E., Ortmann, D., Gentsch, G. E., Mascetti, V. L., Trotter, M. W., Smith, J. C. and Pedersen, R. A. (2015). Brachyury and SMAD signalling collaboratively orchestrate distinct mesoderm and endoderm gene regulatory networks in differentiating human embryonic stem cells. Development 142(12), 2121–2135.

Fathi, A., Eisa-Beygi, S. and Baharvand, H. (2017). Signaling molecules governing pluripotency and early lineage commitments in human pluripotent stem cells. Cell J 19(2), 194–203.

Funa, N. S., Schachter, K. A., Lerdrup, M., Ekberg, J., Hess, K., Dietrich, N., Honore, C., Hansen, K. and Semb, H. (2015). β-catenin regulates primitive streak induction through collaborative interactions with SMAD2/SMAD3 and OCT4. Cell Stem Cell 16(6), 639–652.

Gabut, M., Samavarchi-Tehrani, P., Wang, X., Slobodeniuc, V., O’Hanlon, D., Sung, H. K., Alvarez, M., Talukder, S., Pan, Q., Mazzoni, E. O., Nedelec, S., Wichterle, H., Woltjen, K., Hughes, T. R., Zandstra, P. W., Nagy, A., Wrana, J. L. and Blencowe, B. J. (2011). An alternative splicing switch regulates embryonic stem cell pluripotency and reprogramming. Cell 147(1), 132–146.

Jennings, N. R. (2000). On agent-based software engineering. Artif Intell 117(2), 277–296.

Kalmar, T., Lim, C., Hayward, P., Muñoz-Descalzo, S., Nichols, J., Garcia-Ojalvo, J. and Martinez Arias, A. (2009). Regulated fluctuations in nanog expression mediate cell fate decisions in embryonic stem cells. PLoS Biol 7(7), e1000149.

Kaul, H. (2019). Respiratory healthcare by design: computational approaches bringing respiratory precision and personalised medicine closer to bedside. Morphologie 103(343), 194–202.

Kaul, H., Cui, Z. and Ventikos, Y. (2013). A multi-paradigm modeling framework to simulate dynamic reciprocity in a bioreactor. PLoS One 8(3), e59671.

Kaul, H., Hall, B. K., Newby, C. and Ventikos, Y. (2015). Synergistic activity of polarised osteoblasts inside condensations cause their differentiation. Sci Rep 5, 11838.

Kaul, H. and Ventikos, Y. (2015a). Dynamic reciprocity revisited. J Theor Biol 370, 205–208.

Kaul, H. and Ventikos, Y. (2015b). Investigating biocomplexity through the agent-based paradigm. Brief Bioinform 16(1), 137–152.

Lee, J. J., Talman, L., Peirce, S. M. and Holmes, J. W. (2019). Spatial scaling in multiscale models: methods for coupling agent-based and finite-element models of wound healing. Biomech Model Mechanobiol 18(5), 1297–1309.

Li, X., Upadhyay, A. K., Bullock, A. J., Dicolandrea, T., Xu, J., Binder, R. L., Robinson, M. K., Finlay, D. R., Mills, K. J., Bascom, C. C., Kelling, C. K., Isfort, R. J., Haycock, J. W., MacNeil, S. and Smallwood, R. H. (2013). Skin stem cell hypotheses and long term clone survival explored using agent-based modelling. Sci Rep 3, 1904.

Lin, Y. T., Hufton, P. G., Lee, E. J. and Potoyan, D. A. (2018). A stochastic and dynamical view of pluripotency in mouse embryonic stem cells. PLoS Comput Biol 14(2), e1006000.

Manfrin, A., Tabata, Y., Paquet, E. R., Vuaridel, A. R., Rivest, F. R., Naef, F. and Lutolf, M. P. (2019). Engineered signaling centers for the spatially controlled patterning of human pluripotent stem cells. Nat Methods 16(7), 640–648.

Martin, K. S., Kegelman, C. D., Virgilio, K. M., Passipieri, J. A., Christ, G. J., Blemker, S. S. and Peirce, S. M. (2017). In Silico and in vivo experiments reveal M-CSF injections accelerate regeneration following muscle laceration. Ann Biomed Eng 45(3), 747–760.

Martyn, I., Kanno, T. Y., Ruzo, A., Siggia, E. D. and Brivanlou, A. H. (2018). Self-organization of a human organizer by combined Wnt and Nodal signalling. Nature 558(7708), 132–135.

Muncie, J., Ayad, N., Lakins, J. and Weaver, V. (2020). Mechanics regulate human embryonic stem cell self-organization to specify mesoderm. bioRxiv, doi: https://doi.org/10.1101/2020.02.10.943076.

Naldi, A., Berenguier, D., Fauré, A., Lopez, F., Thieffry, D. and Chaouiya, C. (2009). Logical modelling of regulatory networks with GINsim 2.3. Biosystems 97(2), 134–139.

Ng, H. H. and Surani, M. A. (2011). The transcriptional and signalling networks of pluripotency. Nat Cell Biol 13(5), 490–496.

Okawa, S. and del Sol, A. (2015). A computational strategy for predicting lineage specifiers in stem cell subpopulations. Stem Cell Res 15(2), 427–434.

Pothen, J. J., Poynter, M. E. and Bates, J. H. (2015). A computational model of unresolved allergic inflammation in chronic asthma. Am J Physiol Lung Cell Mol Physiol 308(4), L384–390.

Prochazka, L., Benenson, Y. and Zandstra, P. W. (2017). Synthetic gene circuits and cellular decision-making in human pluripotent stem cells. Curr Opin Syst Biol 5, 93–103.

Raspopovic, J., Marcon, L., Russo, L. and Sharpe, J. (2014). Modeling digits. Digit patterning is controlled by a Bmp-Sox9-Wnt Turing network modulated by morphogen gradients. Science 345(6196), 566–70.

Regier, M. C., Tokar, J. J., Warrick, J. W., Pabon, L., Berthier, E., Beebe, D. J. and Stevens, K. R. (2019). User-defined morphogen patterning for directing human cell fate stratification. Sci Rep 9(1), 6433.

Rosas, F., Mediano, P., Jensen, H., Seth, A., Barrett, A., Carhart-Harris, R. and Bor, D. (2020). Reconciling emergences: an information-theoretic approach to identify causal emergence in multivariate data. arXiv, arXiv:2004.08220v1 [q-bio.NC].

Sato, N., Meijer, L., Skaltsounis, L., Greengard, P. and Brivanlou, A. H. (2004). Maintenance of pluripotency in human and mouse embryonic stem cells through activation of Wnt signaling by a pharmacological GSK-3-specific inhibitor. Nat Med 10(1), 55–63.

Saunders, R., Kaul, H., Berair, R., Gonem, S., Singapuri, A., Sutcliffe, A. J., Chachi, L., Biddle, M. S., Kaur, D., Bourne, M., Pavord, I. D., Wardlaw, A. J., Siddiqui, S. H., Kay, R. A., Brook, B. S., Smallwood, R. H. and Brightling, C. E. (2019). DP2 antagonism reduces airway smooth muscle mass in asthma by decreasing eosinophilia and myofibroblast recruitment. Science Transl Med 11(479), eaao6451.

Saunders, R. M., Kaul, H., Berair, R., Singapuri, A., Chernyaysky, I., Chachi, L., Biddle, M., Sutcliffe, A., Laurencin, M., Bacher, G., Bourne, M., Pavord, I. D., Wardlaw, A., Siddiqui, S., Kay, R., Brook, B. S., Smallwood, R. and Brightling, C. E. (2017). Fevipiprant (qaw039) reduces airway smooth muscle mass in asthma via antagonism of the prostaglandin D2 receptor 2 (dp2). Am J Respir Crit Care Med 195, A4677.

Segovia-Juarez, J. L., Ganguli, S. and Kirschner, D. (2004). Identifying control mechanisms of granuloma formation during M-tuberculosis infection using an agent-based model. J Theor Biol 231(3), 357–376.

Semrau, S. and van Oudenaarden, A. (2015). Studying lineage decision-making in vitro: emerging concepts and novel tools. Annu Rev Cell Dev Biol 31, 317–345.

Smukler, S. R., Runciman, S. B., Xu, S. and van der Kooy, D. (2006). Embryonic stem cells assume a primitive neural stem cell fate in the absence of extrinsic influences. J Cell Biol 172(1), 79–90.

Sokol, S. Y. (2011). Maintaining embryonic stem cell pluripotency with Wnt signaling. Development 138(20), 4341–4350.

Sun, T., Adra, S., Smallwood, R., Holcombe, M. and MacNeil, S. (2009). Exploring hypotheses of the actions of TGF-beta 1 in epidermal wound healing using a 3d computational multiscale model of the human epidermis. PLoS One 4(12), e8515.

Tao, S., McMinn, P., Coakley, S., Holcombe, M., Smallwood, R. and MacNeil, S. (2007). An integrated systems biology approach to understanding the rules of keratinocyte colony formation. J R Soc Interface 4(17), 1077–1092.

Tewary, M., Dziedzicka, D., Ostblom, J., Prochazka, L., Shakiba, N., Heydari, T., Aguilar-Hidalgo, D., Woodford, C., Piccinini, E., Becerra-Alonso, D., Vickers, A., Louis, B., Rahman, N., Danovi, D., Geens, M., Watt, F. M. and Zandstra, P. W. (2019). High-throughput micropatterning platform reveals Nodal-dependent bisection of peri-gastrulation-associated versus preneurulation-associated fate patterning. PLoS Biol 17(10), e3000081.

Tewary, M., Ostblom, J., Prochazka, L., Zulueta-Coarasa, T., Shakiba, N., Fernandez-Gonzalez, R. and Zandstra, P. (2017). A stepwise model of Reaction-Diffusion and Positional-Information governs self-organized human peri-gastrulation-like patterning. Development 144(23), 4298–4312.

Tewary, M., Shakiba, N. and Zandstra, P. W. (2018). Stem cell bioengineering: building from stem cell biology. Nat Rev Genet 19(10), 595–614.

Vahey, M. D. and Fletcher, D. A. (2014). The biology of boundary conditions: cellular reconstitution in one, two, and three dimensions. Curr Opin Cell Biol 26, 60–68.

Wang, Z., Oron, E., Nelson, B., Razis, S. and Ivanova, N. (2012). Distinct lineage specification roles for NANOG, OCT4, and SOX2 in human embryonic stem cells. Cell Stem Cell 10(4), 440–454.

Warmflash, A., Sorre, B., Etoc, F., Siggia, E. D. and Brivanlou, A. H. (2014). A method to recapitulate early embryonic spatial patterning in human embryonic stem cells. Nat Methods 11(8), 847–854.

Wooldridge, M. (1997). Agent-based software engineering. IEE Proceedings-Software Engineering 144(1), 26–37.

Xu, H., Ang, Y. S., Sevilla, A., Lemischka, I. R. and Ma’ayan, A. (2014). Construction and validation of a regulatory network for pluripotency and self-renewal of mouse embryonic stem cells. PLoS Comput Biol 10(8), e1003777.

Xue, X., Sun, Y., Resto-Irizarry, A. M., Yuan, Y., Aw Yong, K. M., Zheng, Y., Weng, S., Shao, Y., Chai, Y., Studer, L. and Fu, J. (2018). Mechanics-guided embryonic patterning of neuroectoderm tissue from human pluripotent stem cells. Nat Mater 17(7), 633–641.

Yachie-Kinoshita, A., Onishi, K., Ostblom, J., Langley, M. A., Posfai, E., Rossant, J. and Zandstra, P. W. (2018). Modeling signaling-dependent pluripotency with Boolean logic to predict cell fate transitions. Mol Syst Biol 14(1), e7952.

Yan, L., Yang, M., Guo, H., Yang, L., Wu, J., Li, R., Liu, P., Lian, Y., Zheng, X., Yan, J., Huang, J., Li, M., Wu, X., Wen, L., Lao, K., Qiao, J. and Tang, F. (2013). Single-cell RNA-Seq profiling of human preimplantation embryos and embryonic stem cells. Nat Struct Mol Biol 20(9), 1131–1139.

